# CARDIAC PIEZO 1 CHANNELS MODULATE ANXIETY

**DOI:** 10.64898/2026.02.18.706706

**Authors:** Jack Summers, Zachary Miklja, Alexander Weaver, Judy Taha, Aiden Yakimchuk, Clayton Kramm, Yong Li, Wael Taha, Luis Afonso, Stephen Farrow, Ryan Woodman, Warren Lockette

## Abstract

Increases in *conscious* cardiac interoception explain the ability of some individuals with anxiety to “feel” their heartbeat without taking their pulse. *Subconscious* cardiac interoception is the detection of heart signals from baroreceptors without subjective awareness. We tested our hypothesis that contrasting sensitivity of PIEZO 1 stretch channels mediates both forms of cardiac interoception and feelings of anxiety. In healthy volunteers, we found conscious cardiac interoception assessed by measuring heartbeat detection accuracy was increased with lower heart rates and greater stroke volumes and cardiac stretch but not associated with anxiety. Disruption of Piezo 1 in cardiac sensory neurons enhanced anxiety in rodents. Accordingly, we used an *ex vivo* assay of PIEZO 1 sensitivity to correlate subconscious cardiac interoception with anxiety in men and women. When compared to healthy individuals with lower PIEZO 1 sensitivity, men and women with *higher* PIEZO 1 activity had marked diminution in state anxiety. Those with lower anxiety also had enhanced variability in their instantaneous baroreceptor sensitivity at rest and reduced cardiac rate pressure products following stress. We propose that a reduction in PIEZO 1 sensitivity causes errors in “predictive coding;” the imbalance between expected and actual heart rate responses to changes in blood pressure leads to anxiety and increased cardiac workloads. We also report testosterone, which is anxiolytic, enhanced, whereas the stress hormone corticosterone, decreased Piezo 1 gene transcription. Selectively enhancing subconscious interoception by increasing PIEZO 1 sensitivity may improve predictive coding, cardiovascular outcomes, and ameliorate the subjective manifestations of anxiety.

## BACKGROUND

Higher heart rates^1^ and lower heart rate variability^2^ are associated with a greater prevalence of anxiety and cardiovascular morbidity.^3^ Although transient fear may be protective for an organism by engaging the fight or flight reflex, men and women with chronic apprehension have persistent activation of their sympathetic nervous system, higher heart rates, greater cardiac work, and increased allostatic load which result in poor cardiovascular health. Conversely, reductions in resting heart rate and increases in heart rate variability correlate with diminished worry and improved cardiovascular outcomes.^1,2,3,4,5^ By measuring increases in heart rate from adrenal catecholamine secretion in stressed laboratory animals, Walter B. Cannon first concluded over a century ago that fear *induces* a catecholamine-dependent increase in heart rate. ^6,7,8^

There is also strong evidence for the *contrary* view that emotional responses such as fear and anxiety *result from* changes in our heart rate. In other words, our heart rates do not accelerate because we are afraid; we experience fear because our heart rates have increased. Our heart does not “skip a beat” because we have anxiety; we have anxiety because our heart skips a beat. This alternative concept, proposed by James and Lange,^9,10^ is similar to the phenomenon of central command in exercise where there is first a rise in heart rate and blood pressure in anticipation of, and prior to, exertion—not just as a result of the metabolic demands of physical work.^11,12^ In the subconscious recognition of a threat, or in anticipation of exertion, there can be unexpected *or* anticipated physiological responses, respectively. Implicit in this theory is that unanticipated increases in heart rate *originate* the emotions of fear and anxiety in response to the subliminal recognition of threats.

We and others have shown there is less anxiety among those individuals with lower resting heart rates; and more importantly, anxiety can be mitigated by *primary* reductions in heart rate.^13,14^ Indeed, there is a reduction in anxiety in patients treated with ivabradine^15^ or beta adrenergic receptor antagonists such as propranolol^16^ and atenolol^17^ which lower heart rates, or following radiofrequency catheter ablation in the atria among those with supraventricular tachycardia.^18,19^ Notably, ivabradine and atenolol have poor penetration across the blood brain barrier—this is suggestive of a peripheral mechanism of action for the anxiolytic effect of these agents.^20^ In other words, these drugs reduce fear instigated by increases in heart rate. Despite the established association between heart rate and anxiety, individuals vary widely in their affective response to changing heart rate. Consequently, the biological basis for contrasting susceptibility to anxiety and the mechanistic link between lowering heart rate, reducing anxiety, and improving cardiovascular health remain unclear.

Interoception describes the processes in which we detect, consciously and subconsciously, many facets of our “*milieu intérieur*” such as heart rate. Differences in conscious cardiac interoception accounts for the ability of some subjects to “feel” every heartbeat without taking their pulse while others are unable to discern any of their heartbeats. The interoceptive sensory receptors that allow us to consciously identify threats to homeostasis, such as increased heart rates (“my heart is pounding”), dyspnea (“I can’t breathe”), and gastric awareness (“I feel queasy”) are not known.^21^ Whereas anxious patients often express their heart is racing, there has been conflicting evidence whether enhanced *conscious* cardiac interoception predisposes individuals to anxiety.^22^ We offer the tantalizing hypothesis that cardiac PIEZO 1 mechanosensitive channels mediate conscious and subconscious cardiac interoception and anxiety by detecting changes in heart rates in response to fluctuations in blood pressure. We also posit that contrasting sensitivities of PIEZO 1 channels are responsible for individual variations in one’s subconscious interoceptive accuracy, and as a result, the susceptibility to generalized anxiety among the population.

Although it is known PIEZO 1 channels in vascular baroreceptors respond to variations in blood pressure,^23^ it is less well appreciated that these mechanosensitive receptors also respond to changes in heart *rate*. It is likely that PIEZO 1 channels of cardiac sensory neurons operate similarly to skeletal muscle where PIEZO 1 channels respond to both the degree of distensibility, and also, the frequency by which these stretch receptors are activated.^24,25,26^ We contend that reduced sensing of the frequency and pressure by which PIEZO 1 channels are distended in cardiac sensory neurons directly contributes to the subjective feelings of generalized anxiety, and individual contrasts in PIEZO 1 sensitivity to changes in heart rate and blood pressure may be responsible for the predisposition for anxiety among some men and women.

We report that one’s ability to reliably sense one’s heartbeat without taking one’s pulse (i.e. conscious interoceptive accuracy) correlated significantly with two indicators of cardiac stretch—heart rate and stroke volume. Enhanced conscious cardiac interoceptive accuracy alone was *not* correlated with state anxiety. Yet we found targeted disruption of Piezo 1 in cardiac sensory tissues arising from the neural crest in rodents increased anxiety as measured by reduced spontaneous exploratory activity and weight gain. We also report the stress hormone corticosterone^27^ diminished Piezo 1 transcription, and testosterone, which is markedly anxiolytic, enhanced Piezo 1 gene expression.^28,29^ The latter finding could explain the lower prevalence of anxiety in men compared to women.

We next used an *ex vivo* assay of PIEZO 1 channels^30,31,32^ to demonstrate *healthy individuals with reduced sensitivity of PIEZO 1 also have higher levels of state anxiety*—the feelings of tension, worry, apprehension and nervousness that fluctuate with the degree of stress. Because PIEZO 1 senses heart rate and blood pressure, we assessed instantaneous baroreceptor sensitivity in men and women with varying degrees of state anxiety. We observed that individuals with lower PIEZO 1 sensitivity had higher state anxiety and greater reductions in the variation of instantaneous baroreceptor sensitivity when stressed. The inability to finely adjust heart rate in response to changes to blood pressure due to low PIEZO 1 sensitivity, i.e. diminished *subconscious* interoceptive accuracy, could disrupt predictive coding which has been shown to correlate with feelings of anxiety.^33,34,35,36^

In predictive coding, the insular cortex matches “top down” (from brain to heart) guidance of heart rate with “bottom up” input from sensory afferent receptors—in this case from PIEZO 1 of cardiac sensory neurons. When cardiac sensory output is diminished, the bottom-up signals from the baroreceptors are “noisier,” i.e. less variable, and the brain struggles to accurately regulate internal states. This increased uncertainty in the insular cortex from mismatched subconscious interoceptive clues by the PIEZO 1 channel could indicate that the *milieu intérieur* is no longer stable—an internal uncertainty viewed as a threat evoking the fight or flight response.^36^

## METHODS

### Recruitment of healthy human subjects

We recruited healthy men and women from our University community to assess the relationship between PIEZO 1 sensitivity, conscious and subconscious cardiac interoception, and anxiety. No subjects had any history of chronic behavioral disorders or acute anxiety that required clinical attention or treatment. Our study was approved by our institutional review board for the Use of Human Subjects in Research, and each volunteer gave informed consent. Subjects were instrumented for real-time continuous recording of cardiac hemodynamics following ten minutes of rest (blood pressure, heart rate, stroke volumes, and interbeat interval) using photoplethysmography (Finapres™). Instantaneous baroreceptor sensitivity was calculated from the average of the naturally occurring fluctuations in heart rate in response to beat-to-beat changes in blood pressure (*vide infra*).

*Assessment of conscious cardiac interoception* was performed with a heartbeat counting task.^37,38^ Supine subjects resting in a quiet room were asked to count the number of heartbeats they experienced *without taking their pulse* over four-time intervals at 35, 45, 60, and again for 30 seconds in random order. Simultaneously, the heart rates of the subjects were recorded with a two-lead electrocardiogram (EKG). Heartbeat detection accuracy was calculated as the ratio of the number of heartbeats a subject experienced compared to the true number of heartbeats recorded from the EKG. Specifically, we used the formula 1/4 Σ [1 – (|EKG measured heartbeats – subjectively reported heartbeats|) ÷ EKG measured heartbeats]. Heartbeat *awareness* was measured on a 10-point Likert scale by asking subjects how often they were aware of their heartbeat.^39^

### Fitness testing

Because fitness training can affect resting heart rate, stroke volume, heartbeat detection accuracy, and susceptibility to anxiety,^40,41,42^ we measured aerobic fitness in subjects. A validated, modified Harvard Step test^43^ was performed in which the subject stepped up and down on a platform (16” in height) at a constant rate of 30 steps per minute (in cadence with a metronome) for 5 minutes or until exhaustion (i.e. when the subject could not maintain this stepping rate for 15 seconds). The subject’s total number of heartbeats were summed between 1 to 1.5 minutes, between 2 to 2.5 minutes, and between 3 to 3.5 minutes. The “fitness index” (FI) considers the increase in heart rate for a set level of exercise, and also, how quickly an individual’s heart rate recovers from that exercise. Higher FI scores represent greater fitness levels. The FI is calculated: FI = (100 x exercise duration) ÷ (2 x [beats_1-1.5_ + beats_2-2.5_ + beats_3-3.5_]) with higher scores indicating greater aerobic fitness.

*Measurement of state anxiety* was assessed using Likert-scale responses to queries from a highly validated pencil and paper questionnaire (State Trait Anxiety-*State* Inventory) modified (mSTAI) to remove three questions that only reflected generalized negative affectivity such as depression.^44,45^ State anxiety is a temporary, situational feeling of worry or tension (like before a test), while trait anxiety is a stable, long-term personality characteristic reflecting a general tendency to be anxious. Subjects also completed the State Trait Anxiety-*Trait* Inventory.

### Determination of cardiac Piezo 1 expression, generation of Piezo 1 conditional knockouts, and measurement of anxiety in rodents

Rodent procedures were performed with the approval of our Institutional Approval for the Use of Laboratory Animals in Research and followed the ARRIVE guidelines protocol.^46^ To determine whether our hypothesis was plausible, we induced conditional disruption of Piezo 1 expression in rodent sensory afferents arising from the heart. Specifically, we created a conditional mouse knockout of Piezo 1 in all adult neural crest derived neurons, which include vagal sensory afferent neurons and sympathetic neurons, based upon the Cre-Lox system and an advillin promoter.^47^ Mice with exons 20-23 of the Piezo 1 gene were flanked by a loxP site for cre recombinase-mediated excision of this floxed region. These mice (Jax Labs, strain #029213, B6.Cg-Piezo1tm2.1Apat/J) were crossed with Advillin-CreERT2 transgenic mice (Jax Labs, strain #032027, AvCre-ERT2) that express a tamoxifen-inducible Cre recombinase directed by the mouse advillin promoter elements that when induced, Cre recombinase activity results in a loss of Piezo 1 activity not only in sensory neurons, but all neurons derived from neural crest tissues including the adrenal medulla.^47^ Conditional knockouts were induced by feeding a diet supplemented with tamoxifen (Inotiv Products,™ TD#130858), *ad libitum*, which provided approximately 80 mg tamoxifen/kg body weight. Control mice were fed tamoxifen in their chow but did not harbor the conditional knockout. Confirmation of Piezo 1 disruption in the heart was confirmed with RT-qPCR using primers spanning exon 19 and exon 22 of the Piezo 1 gene (nt 2764-3414, Forward: 5’-TCT GTG ATG AAC CTG CTG CT and Reverse: 5’-CAG CGT GAG GAA CAG ACA GT with an expected length of 650 bp.^48^

### Hemodynamic measurements in rodents

Blood pressure and heart rates were measured on awake rodents using automated tail cuff plethysmography (Visitech Systems™). Rodents were weighed weekly, and rodent anxiety was assessed by measuring weight over 12 weeks and by measuring spontaneous exploratory activity in an open field during exposure to 7.5% CO_2_ balanced with 92.5% oxygen.^49^

### Ex vivo measurement of human PIEZO 1 sensitivity in men and women

We used measurements of hypotonicity-induced lysis of erythrocytes to assess subconscious interoceptive accuracy sensed by baroreceptors, i.e. PIEZO 1 sensitivity. We performed a modified osmotic fragility test; in response to hypotonicity, erythrocytes swell. PIEZO 1 induces regulatory volume decreases by activating calcium-activated K^+^, Cl^-^, and water efflux.^30,31,32^ Erythrocytes with diminished PIEZO 1 activity have reduced regulatory volume decreases and are more susceptible to lysis. Conversely, red blood cells with enhanced PIEZO 1 sensitivity have greater regulatory volume decreases and are protected against hypotonic lysis. For our assay, 20 µL of whole blood from a heparinized capillary tube obtained from resting subjects was added to a conical tube containing 5 mL isosmotic sodium phosphate buffer (pH = 7.8) diluted to 1:2, i.e., 50% tonicity. After a 20-minute incubation at room temperature, the tubes are centrifuged at 300 × g for 5 min. 150 µL of the supernatant, containing hemoglobin from the lysed cells, is placed into 96 well plates, and absorbances are read at 540 nm. To adjust for any variation in erythrocyte hemoglobin concentration among individuals, values were expressed as absorbance at 50% tonicity against maximum absorbance with complete lysis in 10% tonicity.

### Instantaneous baroreceptor sensitivity

Instantaneous baroreceptor sensitivity was calculated using the sequence method in RStudio. The sequence method is a non-invasive technique that measures baroreceptor sensitivity by identifying spontaneous, consecutive increases or decreases in systolic blood pressure and the corresponding cardiac interbeat interval (IBI) using photoplethysmography. The script identified sections where systolic blood pressure readings and the interbeat interval both increase or decrease together for at least three consecutive heart beats and performed a simple linear regression using the systolic blood pressure as the independent variable and the IBI interval as the dependent variable.^50^ Filters were also included in the script to ensure that sequence detection was limited solely to quantifiable physiological phenomena and not background noise or false readings. A total increase or decrease of ≥1 mmHg was required for each systolic blood pressure sequence identified in addition to an IBI interval filter where each beat in a sequence must be 0.004s ≤ one beat interval ≤ 2s. The script also factored in a delay of one to five beats from the initial identification of an up or down systolic blood pressure sequence to the subsequent corresponding IBI sequence.^51^ After the simple linear regression was completed for each individual sequence identified, the script generated *r*^2^ values for each sequence identified and retained only those that measured ≥0.85 for metadata analyses. A mean, standard deviation, and coefficient of variation was calculated for the instantaneous baroreceptor sensitivity calculated from both up and down sequences.

### Cold Pressor Test

To evaluate stress-induced changes in heart rate variability and instantaneous baroreceptor sensitivity, subjects underwent a standardized cold pressor test. Following a baseline rest period, subjects submerged their non-dominant arm to the wrist in circulating ice water for three minutes. Water temperature was maintained at a constant 0°C via a high-flow circulation pump and confirmed by a digital thermometer prior to each session.

### Assessment of the effect of testosterone on Piezo 1 expression in rodents

Testosterone has marked anxiolytic effects, and the prevalence of anxiety is much greater in women than among men.^52,53,54^ We administered testosterone for one week to male rodents at a dosage which increased plasma levels three fold. Rodents were sacrificed, and Piezo 1 expression in heart tissue was quantified using RT-qPCR.

### Modification of Piezo 1 expression in cardiac sensory neurons

Afferent sensory fibers transmit information from the baroreceptors of the heart through the sympathetic ganglia and also via the dorsal root ganglia and nodose ganglia to the spinal cord and brainstem. The afferent innervation is distinct from efferent innervation but uses the same “highways” for neurotransmission. Afferent cardiac sensory sympathetic neurons, vagal afferent sensory neurons, and adrenal chromaffin cells all arise from neural crest tissue and express Piezo 1. Accordingly, we were able to use PC12 cells to study the effects of 1.0 μM corticosterone (the principal stress hormone in rodents) or 1.0 μM acetylcholine (the principal neurotransmitter for these neurons) on Piezo 1 expression. PC12 cells were incubated with these agents for 24 h and mRNA was isolated and quantified with RT-qPCR.

### Statistical Analysis

Data were analyzed using commercially available software (Prism 10, GraphPad™). Data were examined for normality with a Shapiro-Wilk test for small sample size,^55^ and the robust regression and outlier test (ROUT) was used to ensure there were no significant outliers.^56^ Means for normally distributed categorical data were compared with an unpaired t-test, and Welch’s correction was applied if there were unequal variances; group medians for nonparametric data were compared with a Mann-Whitney test. A p-value (two-tailed) < 0.05 was considered significant.

## RESULTS

To test our hypothesis that stretch receptors in the heart are responsible for one’s ability to feel their heartbeat, i.e. they mediate *conscious* cardiac interoception, we postulated that heartbeat detection accuracy would correlate with cardiac stretch brought about by increased cardiac filling from slower heart rates. The ability to feel one’s heartbeat without taking their pulse in resting, non-distracted subjects was widely distributed within our healthy population cohort of volunteers with a mean value of 79% and a median value of 79% [interquartile range, 54% – 94%]. Even while resting, some subjects could feel *every* heartbeat, while others were barely able to detect their heart rate (Figure 1A). Also, accuracy of heartbeat detection correlated with heartbeat awareness, i.e., subjects that have excellent accuracy of heartbeat detection are more likely to be aware of “feeling” their heartbeat—they had high heartbeat “awareness,” Spearman’s *r* = 0.7507, *p* < 0.0001, *n* = 30. Interestingly, none of the subjects with high heartbeat detection accuracy could describe how they experienced or “felt” their heart beating. Those subjects with lower heart rates permitting higher cardiac filling and larger stroke volumes should have more accurate heartbeat detection than those men and women with higher heart rates and lower stroke volumes. As posited, the accuracy of heartbeat detection was inversely correlated with their heart rate, Spearman’s *r* = -0.4639, *p* = 0.0017, *n* = 43.

**Figure 1.**
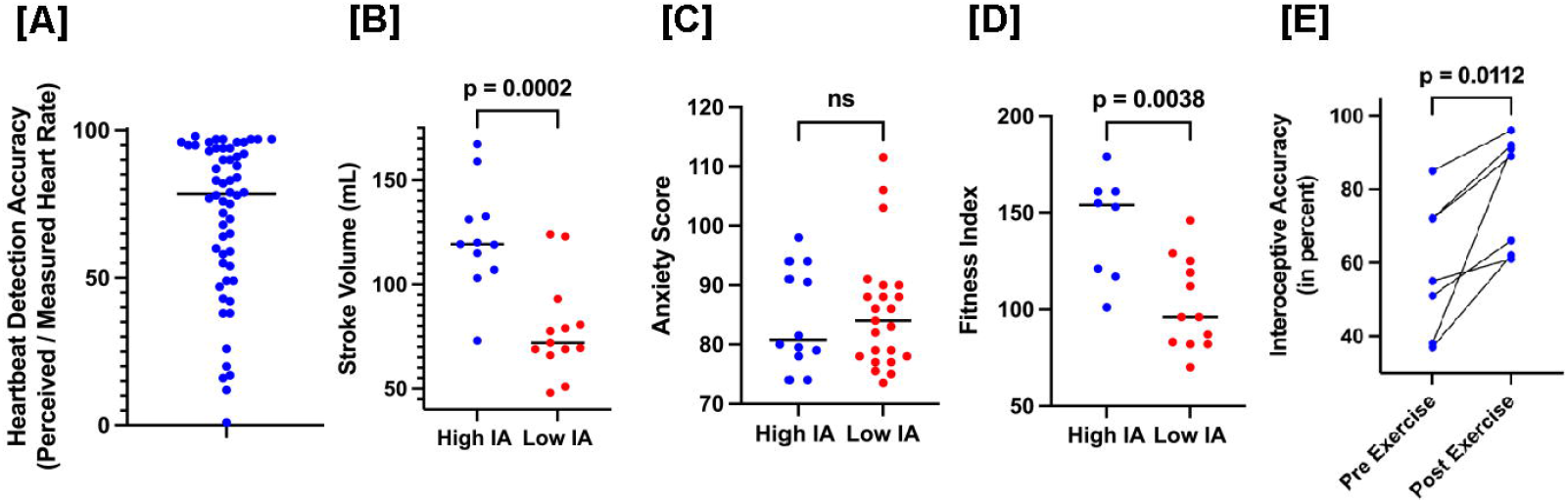
Cardiac stretch allows for conscious cardiac interoception. **(A)** Distribution of heartbeat detection accuracy (“IA”), an assessment of conscious interoception, in resting, non distracted healthy volunteers (n = 56). **(B)** Stroke volume in subjects with high conscious interoceptive accuracy, (above the mean, blue) vs. low (below the mean, red) heartbeat detection accuracy. Horizontal bars represent means. **(C)** Resting state anxiety scores (mSTAI) in high heartbeat detection accuracy vs. low heartbeat detection accuracy subjects. **(D)** Fitness index (modified Harvard step test) in high heartbeat detection accuracy vs low heartbeat detection accuracy among participants who completed fitness testing. **(E)** Effect of acute exercise (modified Harvard step test) on heartbeat detection accuracy.

The accuracy of heartbeat detection was positively associated with stroke volume measured noninvasively with photoplethysmography; those subjects with heartbeat detection accuracy above the mean value had higher stroke volume, whereas men and women with heartbeat detection accuracy below the mean value had reduced stroke volume (values expressed as means in mL ± SEM), 122.4 ± 7.8 vs. 78.6 ± 6.4, p = 0.0002, n = 24 (Figure 1B).

Heartbeat detection accuracy also correlated with the *variation* in an individual’s heart rate (measured as standard deviation of the subject’s continuously measured heart rates at rest, Spearman’s *r* = 0.6736, *p* = 0.0053, and the standard deviation of the real time measurements of systolic blood pressure (Spearman’s *r* = 0.6558, *p* = 0.0071, *n* = 16). In other words, subjects with higher conscious interoceptive accuracy consistently demonstrated more variation in their beat-to-beat heart rates and blood pressures.

If increased awareness of one’s heartbeat or heartbeat variability results in anxiety, then those individuals with high heartbeat detection accuracy, i.e. high conscious cardiac interoception, should be more prone to anxiety. However, scores from subjects in a cohort completing a pen and paper questionnaire, the modified State-Trait Anxiety Inventory (mSTAI),^44,45^ did *not* correlate with a subject’s ability to accurately detect their heartbeat. This vehicle gave good dispersion of state anxiety scores, the median [interquartile range] was 84 [78 – 90]. The ability to detect one’s heartbeat *<u>alone</u>* was *not* associated with a predisposition for anxiety measured by the mSTAI. In healthy subjects who were unstressed and at rest, modified state anxiety scores correlated strongly with trait anxiety scores, *r* = 0.7886, p < 0.0001, n = 19. However, those with strong conscious cardiac interoceptive accuracy (i.e. high heartbeat detection accuracy) scores above or below the mean value did *not* correspond with either high or low state anxiety (values expressed in mSTAI scores ± SEM), 84.5 ± 2.4 vs. 85.6 ± 2.1, p = n.s. (Figure 1C).

Because many of our subjects engaged in regular aerobic exercise training known to diminish state anxiety,^57,58^ we assessed whether fitness levels confounded our measurements of conscious cardiac interoceptive accuracy. Also, many men and women who do not feel their heartbeat at rest can become aware of their heart rate during strenuous exertion.^9^ Because fitness-trained men and women have higher stroke volumes, and because exertion enhances stroke volume,^59^ we expected those with greater fitness would have more accurate heartbeat detection. Aerobic fitness scores from a cohort of subjects performing the Harvard step test correlated with heartbeat detection at rest, Spearman’s *r* = 0.6292, *p* = 0.0314; individuals with high cardiac interoceptive accuracy had much greater fitness scores when compared to individuals with lower heartbeat detection accuracy from (median values expressed in Fitness Index [interquartile range]) 154 [118 – 161] vs. 96 [82 – 123], *p* = 0.0038 (Figure 1D).

If exercise-induced changes in stroke volume are responsible for more accurate heartbeat detection with exertion, then acute exercise should also increase heartbeat detection accuracy. Acute exercise increased heart rates in our subjects by 33% from 61 ± 5 to 91 ± 4, *p* < 0.0001; the exertion-induced increases in heart rates correlated with the resting heart rate, *r* = 0.7650, p = 0.0451, n = 7. More importantly, the accuracy of heartbeat detection immediately following exercise increased from (values expressed as mean ± SEM) from 59% ± 7% to 80% ± 6%, *p* = 0.0112, *n* = 7 (Figure 1E). These results are consistent with the assertion that sensing cardiac stretch is responsible for heartbeat detection accuracy, i.e., conscious cardiac interoception. However, high conscious heartbeat detection accuracy alone is not sufficient to cause anxiety.

Interoception cannot be measured in rodents; however, we were able to assess the effect of conditional disruption of the Piezo 1 gene in sensory afferents on anxiety in mice. We next determined whether ablation of Piezo 1 channels in cardiac sensory afferents which sense stretch of the heart was associated with anxiety. Animals with the conditional knockout in sensory afferent neurons had markedly diminished Piezo 1 expression compared to the control group (values expressed in 2^-Ct^ [interquartile range]) 1.59 × 10^-11^ [9.85 × 10^-12^ – 1.12 × 10^-10^] vs 3.12 × 10^-10^ [3.48 × 10^-11^ – 5.16 × 10^-8^], respectively; *p* = 0.004 (Figure 2A). There were no obvious phenotypic differences in mice with conditional knockout of Piezo 1. Specifically, compared to the control animals fed tamoxifen supplemented diet, there were no differences in awake systolic blood pressure (values expressed in mm Hg ± SEM), 100 ± 3.2 vs. 104 ± 6.7, *p* = n.s., or heart rate (values expressed in beats/min ± SEM), 486 ± 30 vs. 507 ± 20, p = n.s.

**Figure 2.**
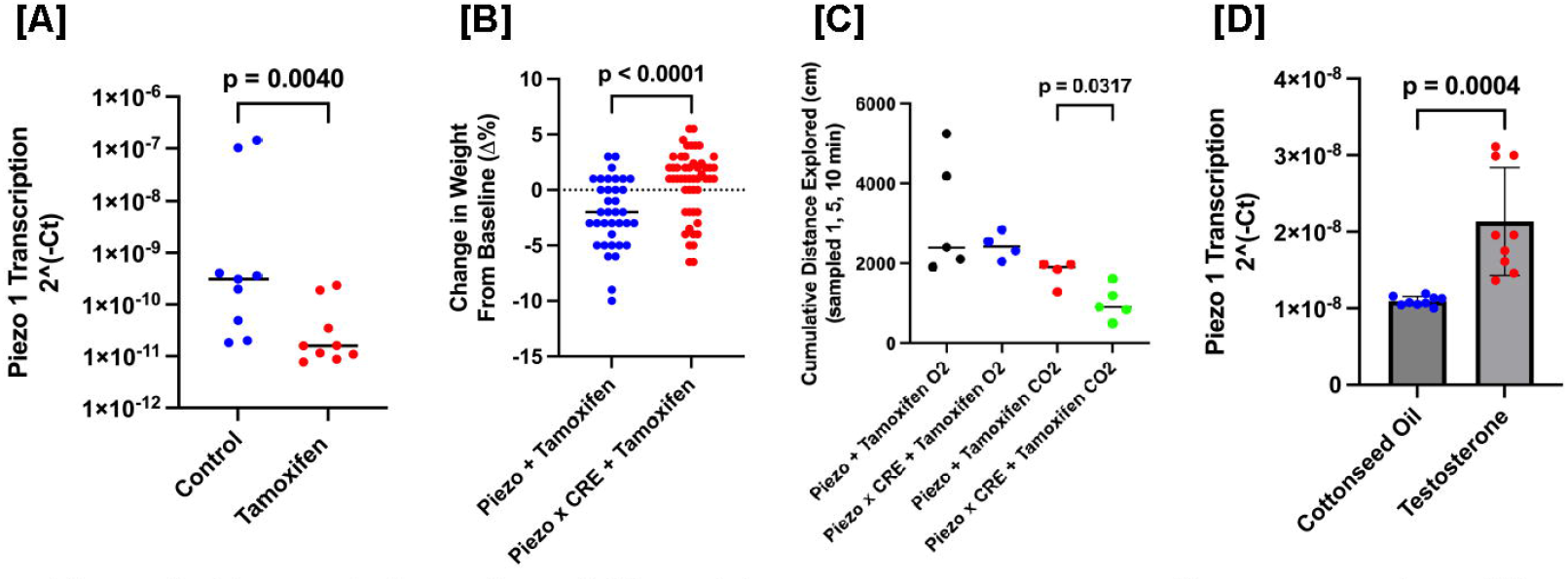
Targeted disruption of Piezo 1 in sensory neurons contributes to anxiety. **(A)** Relative Piezo 1 mRNA expression in heart tissue of conditional knockout mice vs. controls fed tamoxifen. Residual expression reflects Piezo 1 in non-neural crest heart tissue (myocytes/endocardium). **(B)** Percent change in body weight from baseline over the monitoring period in control and knockout mice. **(C)** Anxiety-like behavior measured by cumulative distance explored in an open field during a 7.5% CO_2_ challenge. **(D)** Cardiac Piezo 1 gene transcription in mice supplemented with testosterone (red) was greater when compared to vehicle (cottonseed oil, blue) administered for one week and measured with RT-qPCR.

Unexpectedly, we found when fed tamoxifen, the conditional knockouts *gained significantly more weight* over time when compared to control rodents (values expressed as percent change from baseline weight in percent ± SEM,) -2.14 ± 0.52% vs. +0.57 ± 0.42%, *p* < 0.0001 (Figure 2B). When exposed to 7.5% CO_2_, the knock-out animals also demonstrated less spontaneous exploratory activity in an open field (values measured as centimeters traveled in an open field over 10 min ± SEM) 1016 ± 186 vs. 1773 ±167, *p* = 0.0317 (Figure 2C). Conversely, one week of subcutaneous testosterone administration (which increased steroid levels threefold) significantly enhanced cardiac Piezo1 expression compared to vehicle-treated controls (values expressed as mean 2^-Ct^ ± SEM), 2.1 × 10^-8^ ± 2.4 × 10^-9^ vs 1.1 × 10^-8^ ± 2.0 × 10^-10^, respectively; *p* = 0.0004 (Figure 2D).

PIEZO 1 sensitivity correlates with anxiety in healthy men and women. We used the modified State Trait Anxiety Inventory-State to assess feelings of anxiety in our subjects. To ensure reproducibility of state anxiety testing, fifteen subjects had repeat state anxiety scoring on two occasions separated by at least one week, and the coefficient of variation between these scores was low, 3.3% ± 0.9%. We next measured PIEZO 1 sensitivity in our subjects using an *ex vivo* erythrocyte lysis assay. There was dispersion among our subjects in the degree of lysis of red blood cells, i.e. erythrocyte PIEZO 1 sensitivity, when exposed to gradually increasing levels of hypotonicity (Figure 3A). *Most significantly, the degree of hypotonicity-induced lysis correlated strongly with state anxiety, r* = -0.6495, *p* = 0.0004, *n* = 25 (Figure 3B). Because respiratory acidosis induced by CO_2_ inhalation caused anxiety in our rodents, we also assessed the effect of reducing pH on our *ex vivo* assay of PIEZO 1. Decreasing pH was associated with a marked diminution in the sensitivity of PIEZO 1 activity (Figure 3C); a fall in pH from 7.4 to 7.2 diminished the sensitivity of PIEZO 1 by 14% ± 1.7% but with a wide range of 2.5% to 22.8% among subjects. Erythrocyte PIEZO 1 sensitivity had no effect on conscious heartbeat detection accuracy; i.e. there were no differences in *conscious* interoceptive accuracy between subjects with high vs low PIEZO 1 sensitivity scores, (values expressed in percent heartbeat detection accuracy ± SEM) 75% ± 6% vs. 67% ± 12%, p = n.s. (Figure 3D).

**Figure 3.**
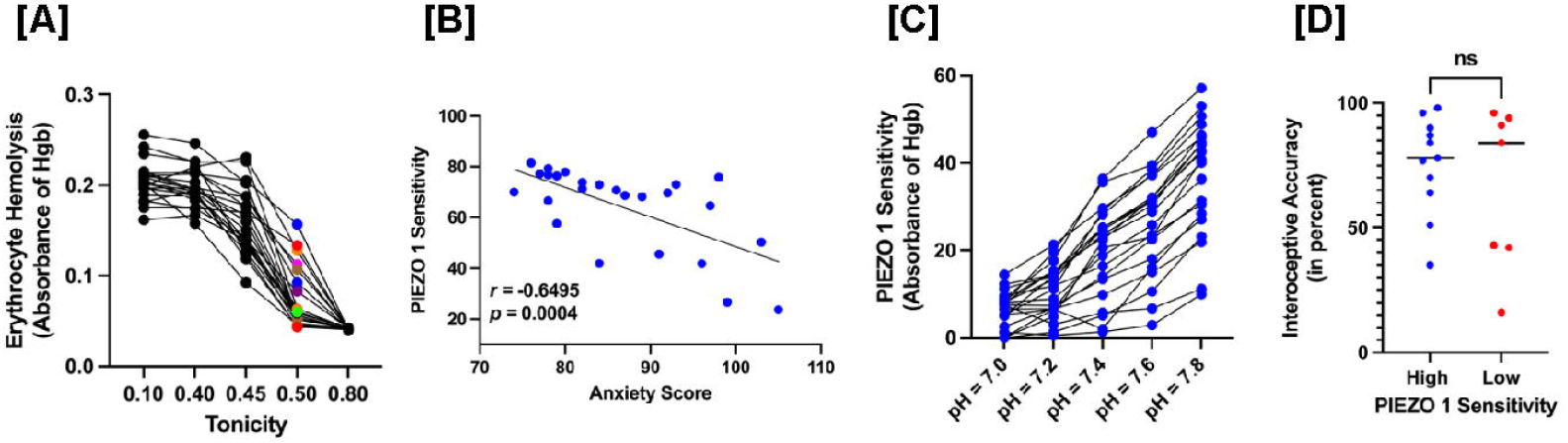
Increases in PIEZO 1 sensitivity correlates with diminished state anxiety. (A) Representative osmotic fragility curves from healthy subjects. Erythrocytes were incubated in buffer of varying tonicity. PIEZO 1 sensitivity is assessed by measuring the protection against hypotonicity-induced erythrocyte lysis. **(B)** Correlation between PlEZO 1 assessed with the erythrocyte lysis assay and state anxiety (mSTAI). The index was calculated as: PIEZO 1 index= 100 x [1 - (A_50_% / A_10_%)], where A_50_% is absorbance at 50% tonicity and Arn% is absorbance at 10% tonicity (maximal lysis). Higher values indicate less lysis (i.e. higher PIEZO 1 sensitivity). **(C)** Effect of extracellular pH on PIEZO 1 sensitivity. There was wide inter-individual variation in pH sensitivity to lysis induced by hypotonicity. **(D)** Conscious cardiac interoception measured by the heartbeat counting task was not different between those with high (blue) or low (red) PIEZO 1 sensitivity.

Increased baroreceptor sensitivity measured as the mean changes in heart rate in response to increases or decreases in blood pressure, is accompanied by enhanced *<u>variation</u>* in baroreceptor responsiveness. Although PIEZO 1 sensitivity did not correlate with the mean values of baroreceptor sensitivity for our subjects, the mean values for instantaneous baroreceptor sensitivity correlated highly with the variability in the measurements of instantaneous baroreceptor sensitivity in each subject, *r* = 0.8062, *p* < 0.0001, *n* = 13 (Figure 4A). Those with the highest PIEZO 1 sensitivity from our *ex vivo* assay demonstrated greater variability in the measurements of instantaneous baroreceptor sensitivity, Spearman’s *r* = 0.7857, *p* = 0.048. In other words, those with high PIEZO 1 activity measured with our PIEZO 1 assay demonstrated greater beat-to-beat variability in their instantaneous baroreceptor sensitivity rather than higher values of instantaneous baroreceptor sensitivity.

**Figure 4.**
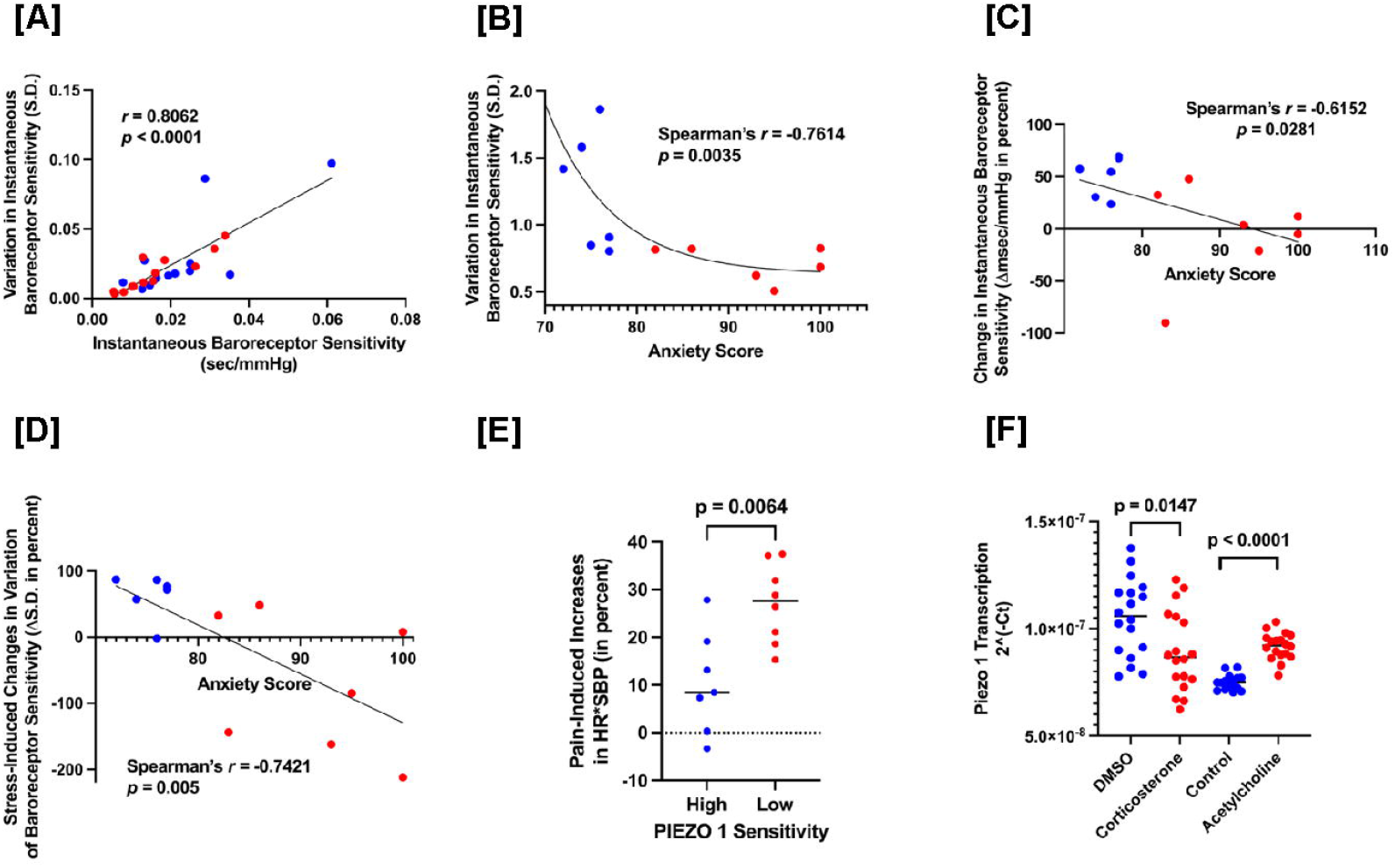
State anxiety correlates with diminished variation in instantaneous baroreceptor sensitivity. **(A)** The standard deviation of values obtained for instantaneous baroreceptor sensitivity from each subject over three minutes was highly correlated with the magnitude of the mean values of each subject for their instantaneous baroreceptor sensitivity. Values were obtained at rest (blue) and during the stress of the cold pressortest (red). **(B)** Subjects with higher anxiety (red), when compared to subjects with less state anxiety, (blue) had lower coefficients of variation (i.e. standard deviation of measurements of instantaneous baroreceptor sensitivity divided by the mean value of the instantaneous baroreceptor sensitivity). **(C)** Subjects with higher anxiety from the pen and paper anxiety inventory had greater reductions in their baroreceptor sensitivity when stressed with the cold pressor test. **(D)** Subjects with higher state anxiety from the inventory also had greater reductions in the standard deviation of their baroreceptor sensitivity induced by the cold pressor test. It should be noted that stress-induced reductions in instantaneous baroreceptor sensitivity and stress-induced reductions in the variability of the instantaneous baroreceptor sensitivity were highly correlated (*r* = -0.8297, *p* = 0.0008). **(E)** Subjects with high PIEZO 1 sensitivity (blue) measured by the erythrocyte assay, when compared to subjects with low PIEZO 1 activity (red), had markedly *reduced* cardiac work in response to the cold pressortest. **(F)** Using PC12 cells as a model of cardiac sensory neurons derived from neural crest tissue, we demonstrated that incubation with the stress hormone corticosterone diminished Piezo 1 expression. Conversely, addition of acetylcholine to mimic a state of high parasympathetic tone (e.g. as seen with fitness training) enhanced Piezo 1 gene transcription.

Consistent with this finding, subjects *with lower state anxiety* measured with the pen and paper anxiety inventory showed much more fluctuations in the beat-to-beat variation in their baroreceptor sensitivity at rest when compared to subjects with higher anxiety, Spearman’s *r* = -0.7614, *p* = 0.0035 (Figure 4B). Most significantly, when stressed with three minutes of the cold pressor test, compared to subjects with lower anxiety, those with higher anxiety responded with much greater reductions in both baroreceptor sensitivity, Spearman’s *r* = - 0.6152, *p* = 0.0281 (Figure 4C), *and* greater reductions in the *variability* measured by the standard deviation of the instantaneous baroreceptor sensitivity over three minutes, Spearman’s *r* = -0.7421, *p* = 0.005. (Figure 4D). *In summary, lower PIEZO 1 sensitivity and higher state anxiety correlate best with <u>lower variability</u> in one’s instantaneous baroreceptor sensitivity*.

### Loss of variability in baroreceptor sensitivity creates more work for the heart

During rest, increases in blood pressure are normally met with reflex decrements in heart rate. However, *stress-induced* increases in blood pressure are not met by corresponding reductions in heart rate, but instead, there is an accompanying increase in heart rate. Unmatched reductions in heart rate following stress induced increases in blood pressure would be expected to increase cardiac workload during stress; indeed, a higher rate pressure product (HR X SBP) denotes greater heart workload. We found that cold pressor test induced changes in cardiac work were much greater in subjects with lower PIEZO 1 sensitivity when compared to subjects with higher PIEZO 1 sensitivity (values expressed in percentage increase in rate pressure product ± SEM), 27.1 ± 2.9 vs. 10.4 ± 4.1, *p* = 0.0064 (Figure 4E). Subjects with low PIEZO 1 sensitivity and anxiety have not only less variability in their instantaneous baroreceptor sensitivity, but also, their hearts work harder when stressed.

Allostatic load is the effort required to maintain fixed (homeostatic) internal set points (e.g. heart rate or blood pressure) in the face of a changing environment (e.g. during psychosocial stress or the cold pressor test). Chronic stress enhances allostatic load through several mechanisms such as enhanced stress hormone production. Conversely, there are several mechanisms to reduce allostatic load such as increasing parasympathetic tone from exercise training.^60^ Having observed correlations between low PIEZO 1 sensitivity, anxiety, and higher cardiac workloads, we assessed whether Piezo 1 expression in cardiac sensory afferents could be modified by autocrine factors associated with increased or diminished allostatic load.

Because cardiac sensory afferent neurons (sympathetic and parasympathetic) and PC12 cells are derived from neural crest cells and express Piezo 1, they can serve as a model to examine control of Piezo 1 expression in response to physiologic changes in these neurons. We first assessed the effect of the rodent stress hormone corticosterone on Piezo 1. Transcription of Piezo 1 was diminished by incubation of neural crest PC12 cells with corticosterone (values expressed as 2^(-Ct)^ ± SEM), 1.05 x 10^-7^ ± 4.26 x 10^-9^ vs. 8.94 x 10^-8^ ± 4.41 x 10^-9^, *p* = 0.0147 (Figure 4F). Next, because exercise training enhances parasympathetic activity with lower heart rates, diminished anxiety, and diminished cardiac workloads, we incubated PC12 cells with acetylcholine. This muscarinic (and nicotinic) agonist markedly enhanced Piezo 1 gene transcription (values expressed as 2^(-Ct)^ ± SEM) from 7.49 x 10^-8^ ± 9.5 x 10^-10^ to 9.18 x 10^-8^ ± 1.47 x 10^-9^, *p* < 0.0001 (Figure 4F). It is suggested that physiological changes that serve to enhance or mitigate chronic stress may do so by mediating corresponding changes in PIEZO 1 gene expression.

## DISCUSSION

The American literary giant Edgar Allan Poe suggested a significant role for cardiac interoception in the generation of anxiety. His famous classic, “*Tell-tale Heart*,” published in 1843, relates the story of a murderer going mad as he hears the beating of his dead victim’s heart buried under the floor boards; his interoception revealed his murderous deed to the authorities—he was driven to admit his crime by the anxiety brought about by hearing *his own* heartbeat. “Villains! I shrieked, dissemble no more! I admit the deed! — tear up the planks! — Here, here! — It is the beating of his hideous heart!” We now report that high conscious cardiac interoception *alone* cannot account for all anxiety.

Interoception arose from the evolutionary advantage of being able to map in real time an organism’s physiological state and allow for immediate corrective action to ensure the fitness of the organism, i.e. to maintain homeostasis. The ability to accurately “feel” one’s heart beat without taking one’s pulse is widely dispersed in the population. During highly strenuous exercise, most individuals can sense their heartbeat without taking their pulse; conversely, some men and women can feel every heartbeat *even while at rest*. We found that increased cardiac stretch brought about by greater cardiac filling from slower heart rates, higher interbeat intervals, and higher stroke volumes in resting subjects strongly correlated with enhanced heartbeat detection accuracy. We also found greater cardiac interoception correlated with fitness training, and heartbeat detection accuracy increased during acute exercise. These findings are consistent with our hypothesis that larger stroke volumes are sensed by stretch of PIEZO 1 channels and associated with greater *conscious* cardiac interoception.

However, if enhanced “feeling” of one’s heartbeats were responsible for feelings of anxiety then a dilemma arises, why are exercise and fitness training correlated with enhanced conscious cardiac interoception but *less* anxiety? PIEZO 1 channels are integral to the baroreceptor-mediated heart rate responses to changes in blood pressure and are also candidates for *subconscious* cardiac interoception. From our data, we suggest anxiety ensues when PIEZO 1 receptors cannot match with fidelity changes in heart rate with physiological fluctuations in blood pressures. In essence, diminution in “predictive coding” brought about by reduced subconscious cardiac interoception, i.e. diminished PIEZO 1 sensitivity of cardiac sensory neurons, is responsible for state anxiety in healthy men and women.

Humans have several homeostatic setpoints that keep our *milieu intérieur* within a narrow, “predicted” range—body temperature (37°C), plasma glucose concentration (72-85 mg/dL), or thirst (plasma osmolality 275-295 mOsm/kg) to name just a few. As an example, the brain maintains a strong prior belief or “set point” for the ideal body temperature that is necessary for survival. An error in predictive coding occurs when sensory inputs from thermoreceptors in the skin or core do not match this expected set point temperature; this prediction error drives autonomic responses such as shivering, sweating, or changes in blood flow which encode behavioral responses. During exercise, baroreceptor mediated increases in heart rate are appropriately matched with elevated blood pressure to maintain the metabolic demands and euphoria ensues. However, when heart rate and blood pressure both increase *unexpectedly* from a stress, a mismatch is perceived between expected and actual heart rate, and anxiety ensues. Alternatively, if diminished detection of changes in heart rate and blood pressure results from reduced PIEZO 1 sensitivity, predictive coding could be threatened, and anxiety could ensue. Several studies of neural networks using functional magnetic imaging and other techniques show that concomitant activation of the insular cortex by baroreceptor activation and emotional responses are coincident with the interpretation of interoceptive clues.^61,62,63,64,65^

The arterial baroreflex is a feedback reflex that mitigates variations in heart rate around an arterial pressure set point to meet the perfusion needs of the organism.^66^ High variability in baroreflex sensitivity is in line with the observation that this physiological mechanism is constantly engaged to match heart rate to systolic pressure changes on a *beat-to-beat* basis. The greater the PIEZO 1 sensitivity, the higher the variability in heart rate that can be sensed by the baroreceptors over time. With diminished PIEZO 1 sensitivity, sensory output on heart rate cannot be matched with fidelity to the expected blood pressure set point, and anxiety ensues. Indeed, this is what we observed. Using an *ex vivo* assay, we first identified individuals with higher or lower PIEZO 1 sensitivity. Higher PIEZO 1 sensitivity in our *ex vivo* assay correlated with greater variability in the instantaneous baroreceptor sensitivity and much less anxiety when compared to subjects with low PIEZO 1 sensitivity. Men and women with higher anxiety had less variation and even greater reductions in instantaneous baroreceptor sensitivity following the stress of the cold pressor test when compared with those subjects manifesting low state anxiety.

It is suggested from our data that in individuals with high PIEZO 1 sensitivity, precise afferent feedback could allow the brain to resolve interoceptive predictions efficiently, confirming homeostasis through subtle corrections. Conversely, in those with low PIEZO 1 sensitivity, the bottom-up signal is blunted or ’noisy.’ This could create a persistent prediction error—a chronic discrepancy between the brain’s regulatory expectations and the ambiguous data received. The brain could interpret this interoceptive uncertainty as a loss of physiological control and trigger a compensatory increase in sympathetic arousal (anxiety) to stabilize the system.

It may be possible to modulate PIEZO 1 sensitivity in men and women. The stress hormone corticosterone diminished Piezo 1 gene transcription, so it is tenable that reductions in stress hormones could enhance Piezo 1 expression. Similarly, acetylcholine enhanced Piezo 1 transcription, and maneuvers which increase vagal tone enhance both the variability and the magnitude of instantaneous baroreceptor sensitivity. Clonidine^67^ and beta-adrenergic receptor antagonists^68^ enhance or unmask vagal tone and enhance the variability of baroreceptor sensitivity. Also, deep breathing^69^ and implanted vagal nerve stimulators^70^ have shown to enhance baroreceptor sensitivity, and they have marked efficacy with anxiolysis.

We note that PIEZO 1 is also expressed in the myocytes and endocardium of the heart^71^ which explains why we did not find complete ablation of Piezo 1 in knockout mice under the advillin promoter. With that in mind, it is key that any action to increase PIEZO 1 sensitivity in cardiac sensory neurons of men and women must be mindful of the effect that intervention could have on myocyte PIEZO 1.

We made several serendipitous observations. During our assessment of PIEZO 1 activity, we found exquisite sensitivity to changes in extracellular pH. CO_2_ inhalation is a frequently used technique to evoke anxiety in clinical studies of anxiolytic agents. The mechanism by which this technique induces anxiety has never been known; we suggest that it may be due to diminution in PIEZO 1 sensitivity brought about by the transient reduction in extracellular pH induced by inhalation of CO_2_.^72,73,74^

We also noted that testosterone enhanced PIEZO 1 sensitivity. Hypogonadal men who complain of a “low T” syndrome find not only enhanced libido with replenishment therapy, but they also report that testosterone has improved their mood. We contend that this effect may not be due to enhanced libido, but instead, from reduced anxiety mediated by up-regulation of PIEZO 1 expression by testosterone. Differences in testosterone-dependent expression of PIEZO 1 levels between men and women could also explain the higher prevalence of anxiety among women when compared to men.

Finally, we were surprised to note that when fed tamoxifen, rodents in which Piezo 1 was conditionally deleted had a marked increase in weight gain when compared to the control mice fed the same tamoxifen supplemented diet. We were unable to determine if this observation were due to anxiety related decrements in spontaneous locomotion and diminished caloric expenditure or decreased appetite. Alternatively, these findings could be due to diminished catecholamine output from the adrenal medulla since that tissue is also derived from the neural crest and would be affected by our conditional knockout of Piezo 1 in this tissue.

PIEZO 1 is expressed in human stomachs, and this mechanosensitive stretch receptor may similarly be responsible for anxiety brought about by gastric interoception (“I feel queasy”). Alternatively, a recent study reported food ingestion stretches the stomach and suppresses ghrelin production; mice lacking Piezo 1 in ghrelin producing cells developed obesity.^75^ It would be interesting to assess gastric interoception^76^ in subjects whose PIEZO 1 sensitivity is known. It is possible that PIEZO 1 may similarly link the higher prevalence of obesity among men and women with diminished satiety and anxiety and heart disease.^77^

We recognize several limitations to our study. For example, we used our erythrocyte assay of PIEZO 1 sensitivity as a proxy for cardiac sensory neurons. The assumption that erythrocyte PIEZO 1 sensitivity reflects cardiac afferent sensory neuron PIEZO 1 has not yet been directly validated. However, this assay has been used successfully to identify individuals with hereditary gain of function or loss of function mutations in PIEZO 1 in the population.^30,31,32^ Also, because erythrocytes do not possess significant transcriptional machinery, red blood cells cannot be used to assess how dynamic changes in the *milieu intérieur* immediately affect PIEZO 1 transcription. Monocytes have transcriptional machinery and express PIEZO 1; we are investigating whether they can be used to assess dynamic changes in PIEZO 1 sensitivity brought about by stress.^78^ Similarly, we note that PC12 cells are of rodent origin. Our work on the effect of stress on neural crest cells await investigation using cells derived from human lines. Indeed, in preliminary studies, we found high levels of PIEZO 1 expression in SH-SY5Y cells which are also derived from a neural crest human metastatic neuroblastoma.^79^

Finally, we recognize that our findings are correlative and do not prove causation; our work is a necessary first step. Prospective studies with interventions that are expected to increase PIEZO 1 sensitivity or expression, e.g. fitness training, testosterone supplementation, etc., need to be conducted to determine if the predicted changes in PIEZO 1 sensitivity (or expression) correlate with expected reductions in anxiety.

The National Institutes of Health has made investigations into interoception a priority area for research whose time has come.^80^ This work is a necessary first step at identifying a potentially new role for the PIEZO 1 channel in sensory neurons—mediating subconscious cardiac interoception and an individual’s predisposition for anxiety. Understanding the biology of PIEZO 1 sensitivity and gene expression in sensory neurons offers a potential roadmap towards treatment of anxiety with peripherally acting anxiolytics that also improve cardiovascular outcomes without the sedation, dependence, and interference with cognition that come from centrally acting anxiolytics. Most importantly, enhancing PIEZO 1 sensitivity in cardiac sensory neurons may reduce cardiac work, diminish allostatic load, improve cardiovascular health outcomes, and make the patient feel better.

